# A Machine Learning One-Class Logistic Regression Model to Predict Stemness in Single Cell Transcriptomics and Spatial Omics Datasets

**DOI:** 10.1101/2023.05.04.539461

**Authors:** Felipe Segato Dezem, Maycon Marção, Bassem Ben-Cheikh, Nadya Nikulina, Ayodele Omotoso, Destiny Burnett, Priscila Coelho, Simone Badal, Judith Hurley, Carmen Gomez, Tien Phan-Everson, Giang Ong, Luciano Martelotto, Zachary R. Lewis, Sophia George, Oliver Braubach, Tathiane Malta, Jasmine Plummer

**Affiliations:** Center for Spatial Omics, St Jude Children’s Research Hospital, Memphis, TN, USA; Department of Developmental Neurobiology, St Jude Children’s Research Hospital, Memphis, TN, USA; Department of Clinical Analysis, Toxicology and Food Sciences, School of Pharmaceutical Sciences of Ribeirao Preto, University of Sao Paulo, SP, Brazil; Akoya Biosciences, The Spatial Biology Company, Marlborough, MA, USA; Department of Obstetrics, Gynecology and Reproductive Sciences, University of Miami Miller School of Medicine, Miami, FL, USA; Sylvester Comprehensive Cancer Center, UHealth Medical Systems, Miami, FL, USA; Nanostring Technologies, Seattle, WA, USA; SAiGENCi, University of Adelaide, Adelaide, Australia; Department of Cellular & Molecular Biology, St Jude Children’s Research Hospital, Memphis, TN, USA; Comprehensive Cancer Center, St Jude Children’s Research Hospital, Memphis, TN, USA

## Abstract

Cell annotation is a crucial methodological component to interpreting single cell and spatial omics data. These approaches are often biased and manually curated. Here we harness an existing stemness model for assessing oncogenic states to transform its application to single cell and spatial omic datasets. This one-class logistic regression machine learning algorithm is used to extract transcriptomic or proteomic features from non-transformed stem cells to identify dedifferentiated cell states. We found this method identifies single cell states in metastatic tumor cell populations without the requirement of cell annotation. Finally these stemness indices are applicable across a variety of spatial transcriptomic and proteomic technologies for the identification of oncogenic cell types in the tumor microenvironment.

**Graphical Abstract:** 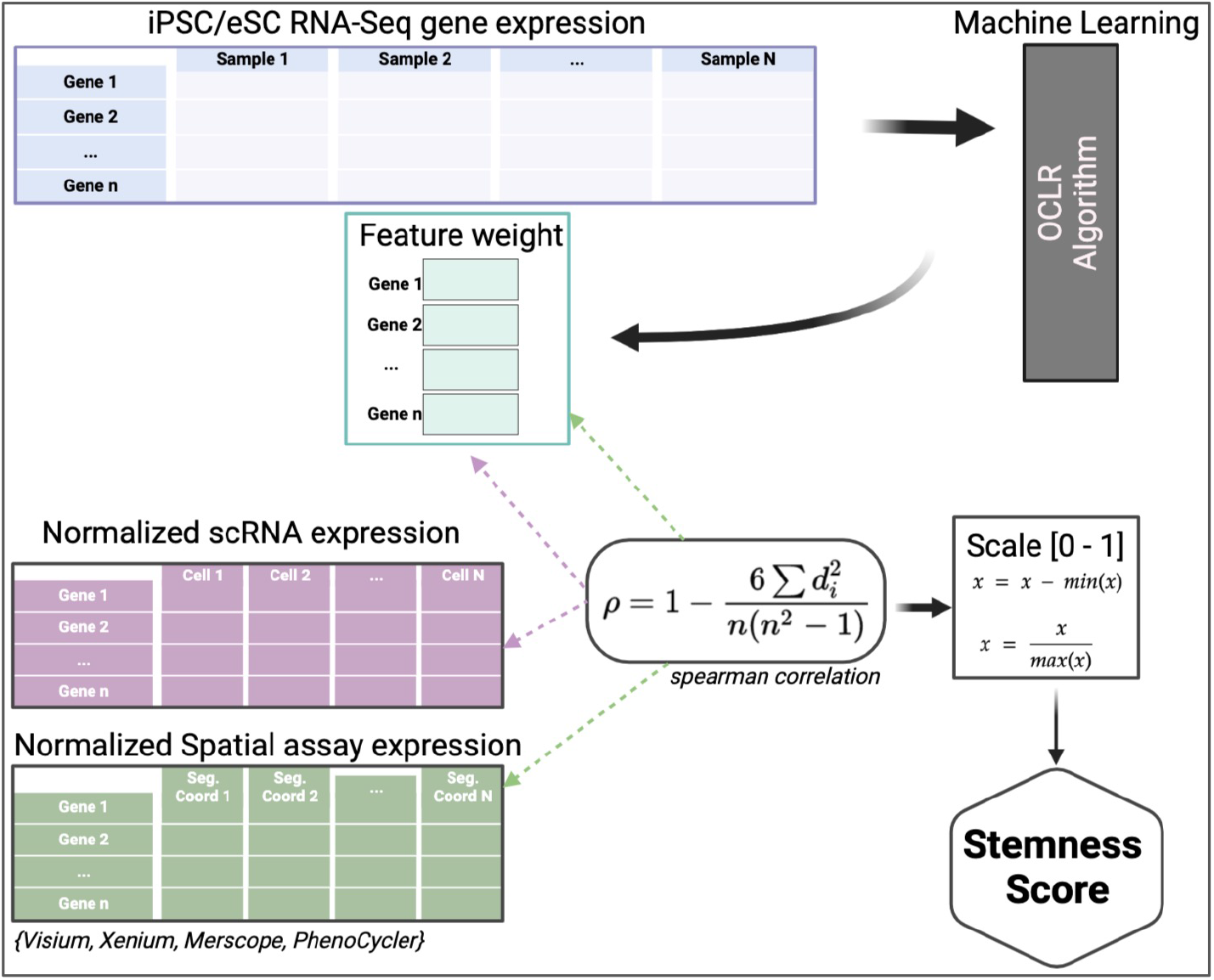

## Introduction

Traditional bulk RNA sequencing provides an average gene expression profile of a population of cells, which may obscure differences among individual cells^1^. In contrast, single-cell RNA sequencing (scRNAseq) has emerged as a powerful method for understanding cellular heterogeneity and allows the measurement of gene expression profiles in individual cells, providing a high-resolution view of transcriptional variability within a population^2^. Advances in scRNAseq technology and computational analysis methods have enabled the identification of rare cell populations, the discovery of novel cell types, and the characterization of cell state transitions during development, disease progression, and treatment response^3^. In this context, scRNAseq has become an essential tool for biological discovery and has numerous applications in diverse fields, such as immunology, neuroscience, and cancer biology.

Despite the demonstrated utility of scRNAseq in aiding discoveries, it has limitations for biological interpretations. scRNAseq data analysis is based on proper cell type annotation. Cell type annotation is used to represent cells as clusters based on their gene expression profiles using unsupervised learning methods^4–6^. These cell clusters are then annotated using these cluster profiles aided by marker gene information. These clustering methods are based on taking previously annotated scRNAseq databases as reference (efforts such as the Human Cell Atlas has been instrumental in aiding this) or using marker genes themselves to annotate cell clusters e.g. CD4 cell clusters are annotated as immune cells, and GFAP positive cells are glial clusters. There are many different subtypes of cell identities for each of these cell types that extend beyond a traditional single marker annotation. The informatic tools and techniques used to identify different cell types, map cellular interactions, and investigate gene expression patterns^7^ are heavily biased and limited to inference for the features/gene expression annotations used in the reference datasets^8,9^. These cell annotations may not reflect the biology seen in the dataset to which it is being applied to. Since each single cell dataset has heterogeneity in the sample processing, sample type and biological treatments, it is difficult to use these methods to match across reference cell annotations or dynamic gene expression profiles. Some methods are developed using the annotated cells from the same dataset to infer the cell types of the remaining but these models require unstructured parameters e.g. number of clusters or are dependent on cell number input requirements^6^.

These problems are further exemplified in the spatial omics field. An explosion of technologies has emerged in spatial omics which are revolutionizing our understanding of complex tissues by providing detailed information on the cellular organization and functioning within intact heterogeneous environments^10^. Spatial omics allows researchers to simultaneously analyze multiple molecular features in the same tissue sample, providing a comprehensive view of cellular interactions, gene expression, and cellular environment. A promising application of spatial omics is the integration of scRNAseq with spatial data to study the heterogeneity of intact tissue. The raw data from spatial omics techniques is often in the form of images or matrices that represent the gene expression across a tissue section^11^. The information captured from spatial omics data is typically processed to generate expression profiles for individual cells, which are then suitable for downstream analysis. The conversion of spatial omics data into single-cell expression profiles is a critical step in the analysis of these data hence the same challenges for scRNAseq analysis exist and are further compounded by the laborious task of cell phenotyping. This is often done by manually assigning cell labels based on the gene marker annotation to the spatial data. These spatial analyses and cell annotation methods share the same challenges as those seen in scRNAseq.

Many Machine Learning (ML) models have been developed to address this problem of large datasets and manual curation. A growing number of deep learning-based methods have been applied to scRNAseq data analyses to enhance the accuracy and efficiency of the analysis and achieve superior performance^12^. These ML models harness generalization and robustness of cell annotations and dataset integration. These algorithms can be used to analyze high-dimensional data and identify patterns that may not be apparent with our present tools. In this study we harness ML algorithms to train and recognize patterns in gene expression data with high accuracy at a much faster rate, hence reducing the time needed to analyze large single cell and spatial datasets to provide new insights into the biology of the tissue being studied.

Annotating cell types in scRNAseq data is important for establishing breast cancer severity and progression based on their tumor microenvironments (TME)^13^. Breast cancer cells that possess stem cell like properties (e.g. self renewal, rapid proliferation) are associated with chemotherapy and radiation resistance, disease recurrence and poorer outcomes^14,15^. ML model generation from bulk tumor RNAseq datasets has been instrumental in identifying breast cancer stemness^16–18^ as the field explores various strategies to target breast cancer stem cells to improve treatment. We apply a bulk RNAseq ML model to single cell and sequencing based, probe based and protein based spatial omics datasets to identify stemness in breast cancer cell types. This study highlights the use of ML in single cell and spatial omic analysis to accelerate our understanding of the complex relationships between stemness, gene expression and tissue morphology.

## Methods

### Data Collection for Reference Sets

The scRNAseq data used for this work was accessed from the Gene ExpressionOmnibus database (GEO; GSE176078 and GSE161529) and ArrayExpress (E-MTAB-6524)^19–21^. We used GSE176078 as our Dataset_1 which contains 26 samples of primary breast cancer, (11 ER+, 5 HER2+, and 10 TNBC) with major and minor cell types already labeled by the authors. Another cohort GSE161529 within 32 samples of primary breast cancer (17 ER+, 6 HER2+, 4 BRCA1 pre-neoblastic, 4 TNBC, 4 TNBC/BRCA1, and 1 PR+) was used as Dataset_2 to validate our main finds in breast cancer cells on stemness context. We then used an induced pluripotent stem cells dataset (E-MTAB-6524) to validate our stemness model in scRNAseq data.

Spatial transcriptomic datasets were retrieved from publicly available datasets from the Vizgen company website (MERSCOPE)^22^, the breast cancer study previously^19,23^. The spatial proteomic datasets were generated by Akoya Biosciences and Nanostring Technologies. The Phenocycler Fusion dataset was run on a FFPE breast cancer sample from the biorepository at University of Miami Health Science Systems. This tumor underwent pathological review and sections were sent to Akoya Biosciences for processing. qpTIFF files were generated and rendered by the company. All files were processed based on 10X Genomics and Akoya’s instructions (see below) and used as input for the stemness model. These processed datasets were in a similar format for gene counts as single cell data and used in the same format as scRNAseq for stemness prediction modeling. The CosMX proteomic dataset was generated and provided by Nanostring Technologies.

### Single cell processing and analysis

Samples from both datasets were processed with the same settings using Seurat v4.0 for QC, filtering, normalization, clustering and visualization. The filtering parameters were as follows: nFeature_RNA > 200 < 5000, percent.mt < 10; nCount_RNA > 200. The cells that have mitochondrial genes greater than 10% or have fewer than 200 detected genes were filtered out. A scale factor of 10,000 was used to normalize all the remaining cells. To correct for the batch effect between different samples, the reciprocal principal component analysis (RPCA) method in the Seurat package was applied to integrate the complete data set. The genes enriched in each cluster were identified using FindAllMarkers function in Seurat. It applies a Wilcoxon Rank Sum test and then performs multiple test corrections using the Bonferroni method. The multiple-test corrected P < 0.05 was used as a cut-off for significance. Samples were normalized using the following settings: normalization.method = “LogNormalize”; scale.factor = 10000>. The remaining Seurat parameters were default with the exception of: FindVariableFeatures:-selection.method = ‘vst’, nfeatures = 3000; RunPCA-features = 3000 VariableFeatures; FindNeighbors-reduction = ‘pca’, dims = 1:20 (20 PCs); FindClusters: resolution 0.5; RunUMAP: reduction = ‘pca’, dims = 1:20. For additional processing and graphical representation the following R packages were used in sc-type v1.0, dplyr v1.1.1, and ggplot2 v3.4.2.

### Spatial data analysis

#### Visium dataset

Visium breast cancer data from 10X genomics was mapped and demultiplexed using 10x SpaceRanger as per company default parameters. Processing included transforming to read counts, overlaying expression data with H&E tissue images and unsupervised clustering. Specifically, spots classified as artifact and spots with count = 0 were removed. Seurat version 5 was used to perform data normalization with the method SCT transform^24^, and data clustering using the Louvain algorithm with multi-level refinement and resolution = 1. For the stemness prediction, we used the normalized expression matrix to calculate spearman correlation with the feature weights from the OCLR model matching genes and scaled to 0-1 format. We then transferred the scores to the Visium seurat object for visualization.

#### Xenium dataset

This dataset included RNA reads, images, and cell segmentation were directly downloaded from 10X website and run using their recommended Explorer pipeline.Molecules with count = 0 were removed and the same analysis routine described previously for Visium data was employed. We then performed cell-type deconvolution using the RCTD method^25^ to predict each cell point present in the xenium dataset using the major cell type annotations from the scRNAseq dataset.

#### Vizgen dataset

Output files from Vizgen dataset were processed as follows: Molecules with count = 0 were removed, SCT was used to transform and normalize the data, with a clip range of [-10,10]. For cell clustering, SCT filter was used at a lower resolution of 0.3.

#### PhenoCycler Fusion dataset

QuPath v0.4 was used to process the 66-channel QPTIFF image generated from the PhenoCycler instrument. Briefly, we employed the StarDist (arXiv:1806.03535) algorithm for cell segmentation using the DAPI channel and exported the measurements as a text file. We selected measurements in cells and mean intensity of captured signal. The DAPI channel was removed for further analysis. To create a Seurat object, we created a sparse matrix with each channel as rows and each segmented cell as columns, and a metadata matrix with centroid coordinates for each cell. Centroids coordinates were used for visualization. In order to predict stemness score for each cell, we created a new model using 28 genes that intersected the panel used in this assay and genes found in the bulk RNAseq data used to train the OCLR model and employed the same strategy to transfer the scores back to the Seurat object.

#### CosMX dataset

The Nanostring dataset was segmented and generated by Nanostring Technologies. The input file was provided as a cell count matrix as per their published pipelines. Seurat object was created using the same method used for the Phenocycler dataset.

### Stemness prediction modeling

Building on existing methods from bulk RNAseq^16^, we extended this iPS/ES model for utilization in single cell RNAseq data. The model parameters were based on the total content of 12 922 genes. The iPS dataset (20 000 cells)^19–21^ was used to validate the iPS/ES bulk model in single cell data. Similarly, this model was tested using two large published breast cancer datasets of 80 000 and 180 000 cells respectively. These three datasets were combined as Seurat objects for normalization and extraction of their gene expression. The stemness model was applied to these single cell datasets using a spearman correlation value of both genes weights in the model and gene expression values on the datasets for each cell in the single cell matrix. We scaled all correlations within 0 to 1 calling these scores to create the stemness index. These individual Seurat objects were used for the next stage of analysis where they were used as input into the Seurat clusterization pipeline.

### Performance of Single Cell Stemness Model

To determine the reliability of the stemness model in single cell data, we performed bootstrapping on both high and low quality samples to test the consistency of the model’s predictions under varying conditions. We compared cell type annotations from dataset 1 and dataset 2 by identifying the most highly expressed cluster markers in dataset 1 and intersecting them with cluster markers in dataset 2. For the major cell type class described in dataset 1 paper^19^, we selected the top 30 highly expressed cell type markers from and intersected them with cluster markers in dataset 2 that had an AvgFoldChange > |0.5|.

Similarly, for minor cell types, we chose the top 10 markers and intersected them with cluster markers in dataset 2 that had an AvgFoldChange > |0.5|. Finally, we used the package sc-type to assign cell types described in dataset 1 to dataset 2. Differential expression analyses were performed to detect marker genes for different cell clusters. The default parameters on Seurat FindMarkers were used for the input for differential expression analysis(pvalue <0.05; FoldChange > |0.5|). Differential expression was performed against the top stemness cells vs bottom stemness cells based on their quantiles (25 percentile vs 75 percentile) within each cell type for both dataset_1 and dataset_2.GO pathway enrichment analyses was conducted using the clusterProfiler package v3.16 ^26^.

### Data visualization

All plotting functions used in this study were included in Seurat v4,Seurat v5, ggplot2, RColorBrewer ^27^ and plotly ^28^ packages.

## Results

We calculated a stemness index for every cell in two independent breast cancer scRNAseq datasets by correlating the gene weight derived from the one-class model trained on bulk RNASeq iPSC/ESC cell lines with the normalized gene expression from the scRNAseq datasets and scaling that metric to [0-1] format. We are able to visualize every stemness index in UMAP formats (Fig 1B, 1C) and based on top 75% and bottom 25% of scoring stemness cells, we calculated that 20,171 of 80,682 total cells in Dataset 1 present a high stemness phenotype, compared to 45,079 of 180,315 of cells in Dataset 2. In the opposite stemness spectrum, 20,171 of cells in Dataset 1 present a low stemness score, compared to 45,079 of cells in Dataset 2.

**Figure 1.**
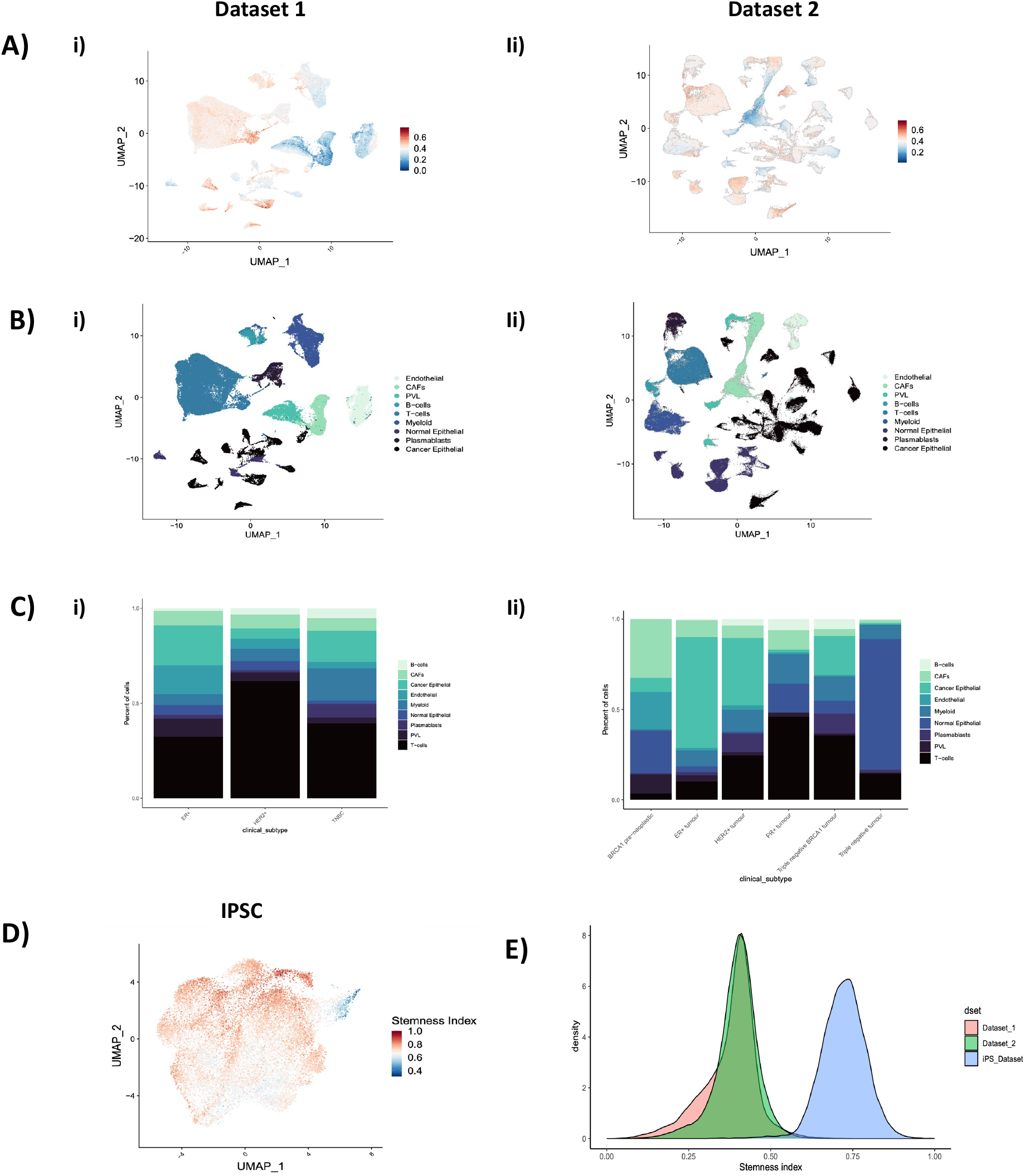
Stemness model using single cell datasets. A) UMAP plot of two independent single cell breast cancer datasets (i, ii). Gradient (blue to red) indicates low to high stemness. B) Cell clustering of major cell types across each dataset. Colors denote the following cell types seen endothelial, cancer associated fibroblasts (CAFs) B cell, T cells, myeloid, normal epithelial, plasmablasts and cancer epithelial cells (from lightest to darkest blue) in dataset 1 (i) and dataset (ii). C) Cell proportion of major cell type clusterings in each breast cancer histotype in both dataset 1 (i) and dataset 2 (ii). Colors align with cell type identification. D) UMAP of iPS single cell dataset used for stemness model. Gradient (blue to red) indicates low to high stemness. E) Stemness index distribution of single cell iPS (blue) compared to breast cancer cells in dataset 1(pink) and dataset 2(green).

We present both datasets using the same cell annotation and we show the proportion of cells per clinical classification in Dataset 1, where samples are divided into ER+, HER2+ and Triple-negative breast cancer (TNBC). These categories are also present in Dataset 2, with the addition of BRCA1 pre-neoplastic, PR+ and TNBC+BRCA1 cells(Fig 1F, 1G). In order to test the biological significance of the stemness score, we applied the same model on a iPS scRNAseq dataset^21^, in a total of 21,599 cells the median stemness score was 0.724 compared to 0.394 and 0.402 in datasets 1 and 2 respectively(Fig 1H).

We associated three different cell types with high or low stemness profiles: i) cycling T-Cells [n=1442 (1.78%) Dataset 1; n=13760 (7.63%) Dataset 2], ii) cancer cycling [n=2631 (3.26%) Dataset 1; n=12125 (6.72%) Dataset 2] and iii) cancer basal cells (n=2877 (3.56%) Dataset 1; n=20156 (11.1%) Dataset 2)(Fig 2A, 2B). In Dataset 1, we observe clusters where cancer cycling and cancer basal cells are clustered together. Cycling T-cells were associated mostly having higher stemness profiles than average in both datasets (Fig 2A, 2C). In Dataset 2, cycling T-cells were mostly clustered with cancer basal cells and more sparsely separated compared to Dataset 1, but with a similar overall stemnes index profile (Fig 2B, 2D). We were able to see this pattern in both breast cancer datasets.

**Figure 2.**
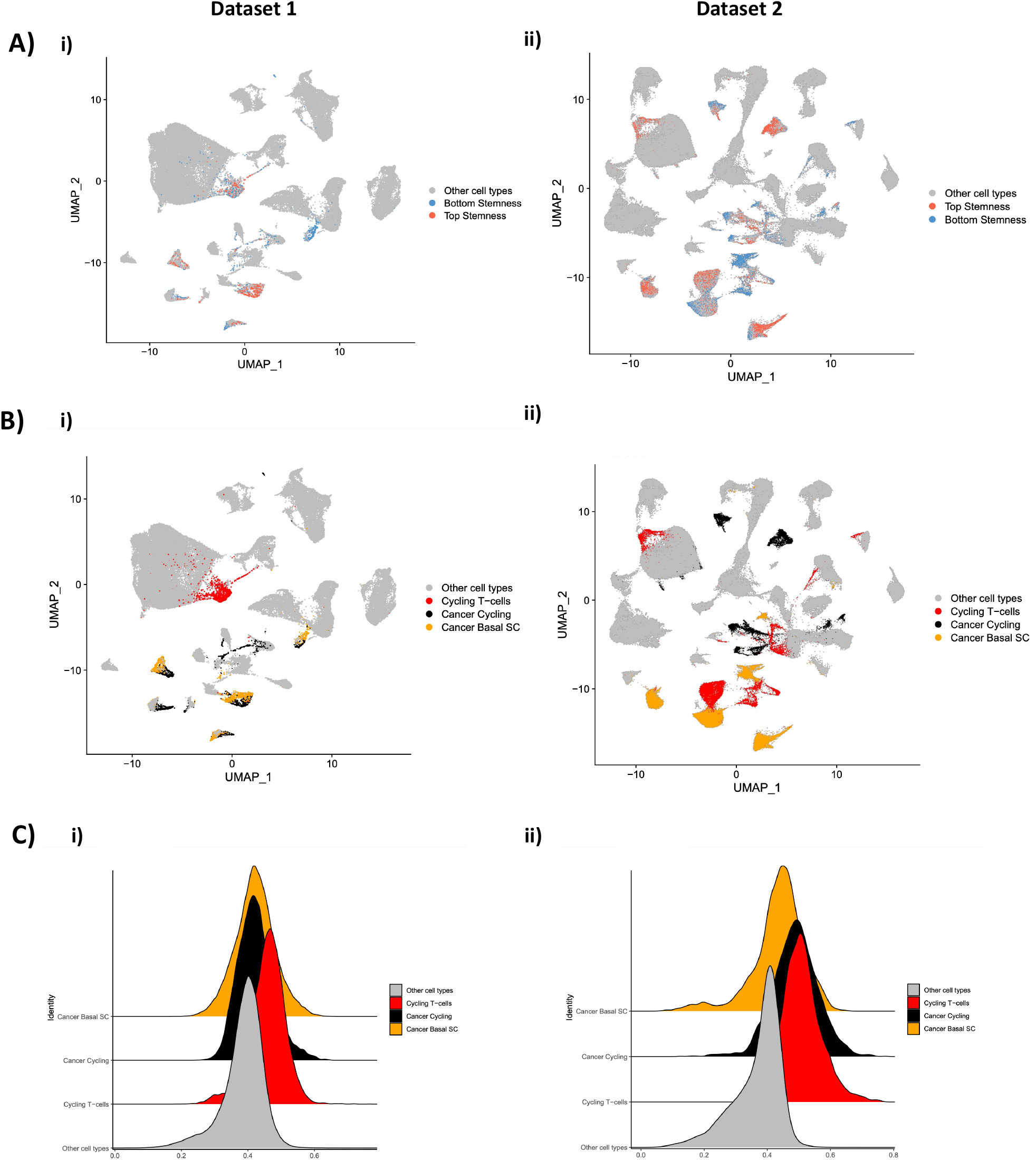
A) UMAP plots of stemness model from two breast cancer single cell datasets (i, ii) of the top (red) and bottom (blue) stemness cells. B) Cancer basal (yellow), cancer cycling (black), and cycling t-cells (red) showing cell types overlapping most top stemness cells in both datasets (i, ii). C) Stemness distribution in dataset 1 (i) and dataset 2 (ii) of breast cancer single cells with the highest stemness seen in cancer basal (yellow), cancer cycling (black) and cycling t-cells (red) compared to all other cells for both datasets (gray).

From the selected cell types we conducted differential gene expression analysis on both datasets. From dataset 1, cancer basal, cancer cycling and cycling T cells have 685, 393 and 230 DEGs respectively. From dataset 2, cancer basal, cancer cycling and cycling T cell had 525, 528 and 632 DEGs respectively. We analyzed the gene expression differences between the selected cell types in both datasets. The groups were separated based on their stemness scores, with one group having higher scores and the other having lower scores. To ensure equal group sizes, we retained the same number of cells in both groups. For cancer basal in dataset 1, we identified 336 genes highly expressed and presenting high stemness scores, among the top expressed genes with high stemness is LDHB (Fig 3A), which plays a role in cancer cells to use lactate more often than glucose that leads to cell proliferation ^29^. For dataset 2, 311 highly expressed genes were identified differentiating top and bottom stemness cells. One of the genes with higher log2 Fold Change is CXCL1 (Fig 3B) and was shown in previous studies^30^ to stimulate invasion and migration in ER negative breast cancer cells. We found 104 genes in common in both datasets, 30.9% and 33.4% of dataset 1 and dataset 2 respectively. We performed GO enrichment analysis using these gene lists as input for both datasets, in dataset 1, (n=72) terms were enriched using the Biological Process ontologies. For dataset 2, that number was (n=91). The number of overlapped terms was (n=17) (Fig 3C). To add another level of confidence that the results seen in dataset 1 and 2 were similar, we performed a semantic similarity analysis^31^, which takes the non-overlapping genes in both datasets as input and computes a similarity score based on the same gene ontology. To establish if the DEGs from dataset 1 and 2 were similar beyond their pathway enrichment overlap, we conducted a gene similarity analysis. In this analysis we took the dataset 1 DEGs and dataset 2 DEGs that define stemness to calculate the similarity index ^31^ to take into account confounding variables in our analysis (Supp. Fig 2E). Our results revealed that the similarity scores between the high stemness gene lists were higher across datasets than the regular clusters.

**Figure 3.**
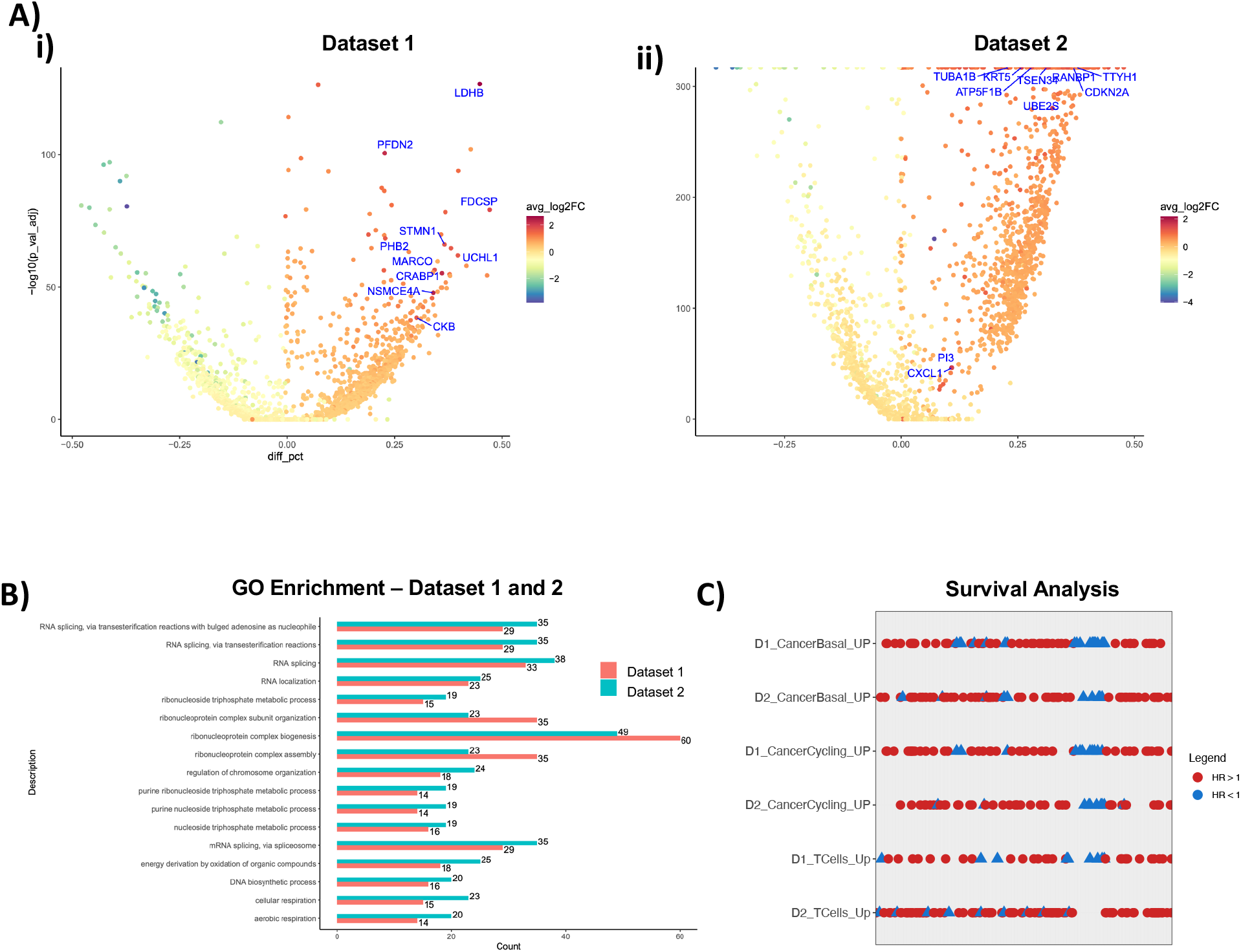
Gene expression analysis of stemness model A) Volcano plots of top and bottom stemness cancer basal cells from dataset 1 (i) and dataset 2 (ii) respectively. Gradient depicts negative logFC (blue) to positive logFC (red). Y axis represents the statistical significance and X axis the different percentage of cells expressing a given gene. Positive different percentage means that top stemness cells have more cells expressing that gene. B) GO enrichment analysis of the top regulated genes in cancer basal top stemness cells for dataset 1 (red) and dataset 2 (blue). The number of DEG is plotted on the X axis and GO categories the genes are represented in are plotted on the y axis. C) Survival analysis from TCGA breast cancer cohort of up regulated genes on top stemness cell type across datasets 1 and 2. Rows represent different gene lists according to dataset and cell type. Hazard ratio > 1 in blue, < 1 in red (p-value < 0.05).

To understand the biological and clinical relevance of these genes identified in the differential expression analysis of high stemness scoring cells, we used gene expression and clinical information data from breast cancer tumors bulk RNA-seq (n=902) from TCGA (The Cancer Genome Atlas)^32^. To correlate gene expression with survival status, we performed a cox regression analysis for each gene with overall survival up to 60 months. We identified 94 genes (27.9%) with a significant (p < 0.05) correlation with poor survival in cancer basal cells for dataset 1, and 70 genes (22.5%) for dataset 2. In cancer cycling cells, we found 51 genes (21.4% and 19.5%) correlating with poor survival in both datasets (Fig 3D) and in cycling T-cells we found 37 genes (19.4%) that correlates with poor survival in dataset 1 and 94 genes (21.5%) in dataset 2.

We sought to apply this single cell derived stemness model into spatial omics (transcriptomic and proteomics) datasets. Breast cancer tumor samples profiled across five different spatial omics assays were analyzed: one sequencing-based, spot based Mwhole-transcriptomics assay; two high plex in situ probe-based imaging platforms and two high plex protein-based platforms. The spatial whole transcriptomics assay (10X Visium) used spatially barcoded RNA sequencing to generate high-resolution maps of gene expression patterns in intact tissue sections. We obtained 17 distinct clusters from 4665 spots and 47774 features (Fig 4A). We used 12,342 features overlapping with the original model to generate new weights to perform correlation and generate stemness scores for each spot. The average stemness score of 0.52 for all spots across the whole sample. For spots demarcated by pathologist classification as invasive cancer, the ML model had the highest average of all classified spots presented of 0.55.

**Figure 4.**
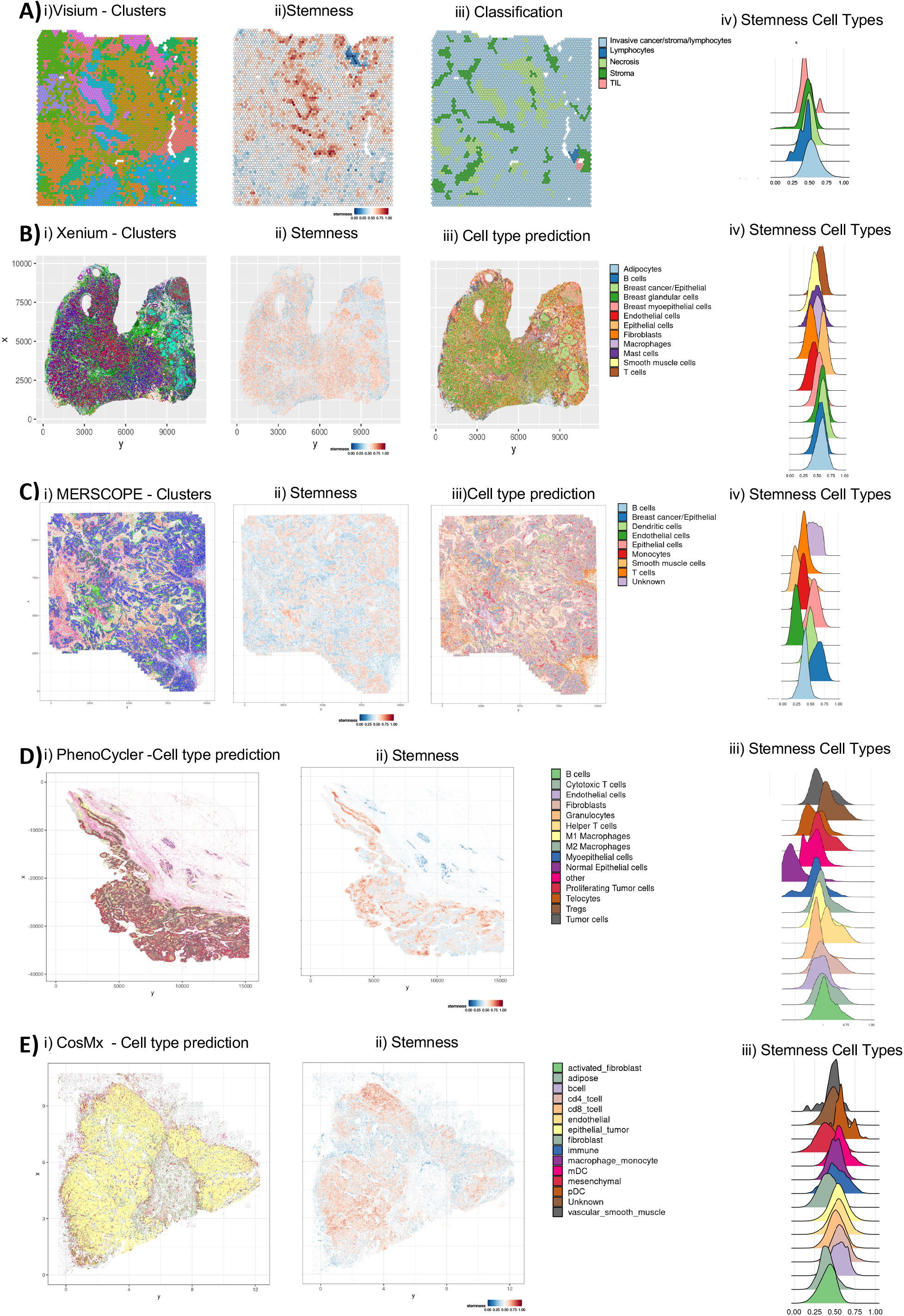
Spatial analysis of stemness model A) Each spot from the section placed on Visium barcoded array is represented with i) a cluster projection ii) stemness score pathology classification for each spot of the Visium sample and distribution plots of stemness score across classified regions (A). Cluster projection, stemness score, cell type prediction for each segmented cell’s centroid location and distribution plot of stemness score per annotated cell types of Xenium sample (B).Cluster projection, stemness score, cell type annotation for each segmented cell’s centroid location and distribution plot of stemness score per annotated cell types of the MERSCOPE sample (C). Cell type annotation, stemness score for each segmented cell’s centroid and distribution plot of stemness score per annotated cell type of the PhenoCycler sample.

We next determined if the ML model was applicable to RNA probe based assays that are not the entire transcriptome. Using a publicly available dataset, we obtained 27 clusters from 889,765 cells profiled from a breast cancer panel (280 genes) on a high plex in situ imaging Xenium platform (Fig 4B). We predicted stemness scores for each segmented cell as described in the methods section. The average stemness score was 0.546. We used the dataset 1 scRNA data to predict cell types based on cell type scoring using the biomarkers present in the antibody panel and the corresponding cell type provided by the company. We created a scoring approach to score each cluster based on the expression of these markers, labeled by the highest scoring cell type.To interrogate the reliability and reproducibility of the ML stemness model across platform types,we tested this model in a publicly obtained dataset from another in situ probe based imaging platform. From the downloaded breast cancer sample from the MERSCOPE Immuno-oncology Data Release, we obtained 18 clusters from 710,073 cells profiled with 500 genes (Fig 4C). The same reference was used to score clusters and assign cell type annotation as described in the Xenium analysis. The average stemness score was 0.474 for all cells, and the two highest scoring cell populations were breast cancer cells and epithelial cells and the lowest cell population stemness score was in fibroblast cells (Fig 4C).

The ML model showed robustness across three distinct spatial transcriptomics methodologies. We next tested its application to high plex proteomic based imaging platforms. A breast cancer sample was analyzed using an immuno-oncology specific 45 plex antibody panel(Discovery Panel, Akoya) and 62 plex panel (Nanostring). From the Akoya sample, 1,040,049 cells were seen across 56 clusters. Based on marker correlation, we annotated 14 different cell types (e.g. cycling cancer cells), similarly seen in both the single cell and spatial transcriptomic data. The average stemness score was 0.516 and the highest scoring cell populations were tumor cells (0.52), Tregs (0.604), proliferating tumor cells (0.525) and helper T cells (0.603). Consistently, normal epithelial cells had the lowest score of all cell types, with an average of 0.246 (Fig 4D). For the CosMx sample, 472,763 cells were identified across 14 cell types, the average stemness score was 0.51 and the highest stemness scoring cell types were epithelial tumor (0.542) and endothelial (0.547). Overall, the average stemness score for all different spatial omics assays were similar, and the cell types presenting a higher stemness profile were also similar.

## Discussion

Our study demonstrates the utility of ML modeling for agnostic cell type annotation in both scRNAseq and spatial omic data^33^. This ML model overcomes the computational hurdles and challenges of analyzing millions of cells against hundreds of features^11^. An advantage to this ML model for scRNAseq and spatial omics analysis also include faster processing time than laborious manual cell phenotyping which is exponentially longer in spatial omics analysis.We demonstrate its robustness across a variety of spatial omics technologies including, sequencing based, RNA/in situ probe based and protein based platforms. Using our stemness scRNAseq and spatial model approach to breast cancer, we identified cell types within existing cell clusters attributable to cancer stemness which were not previously described. This ML approach can identify scRNAseq features that are predictive of clinical outcome, which can help in patient stratification and personalized medicine.

Breast cancer tumor biology is concordant with the stemness predictions of the model^15,34^ such that cell types known to be involved in more aggressive disease have increased stemness (e.g. proliferating tumor cells). Similarly, cells that are responsible for effective immune response (e.g. helper T cells) are decreased in tumors with high stem. This method also addresses the problems of proper cell annotation such that cells regardless of their annotated cell identity can be properly identified based on their level of stemness.Normal epithelial cells without annotation were identified with no levels of stem, while proliferating tumor cells had high levels of stemness. This model can be more broadly applied to various diseases, treatment conditions and genetic differences compared to existing manual cell curation methods.

The interaction of breast cancer cells with their environment relies on communication between local cues, cancer cells, cancer stem cells, immune cells and stromal cells within the tissue^14^. This TME has clearly been linked to prognosis, recurrence, treatment resistance and outcome^14^. For example, breast cancer treatment targets the proliferative advantage breast cancer cells have compared to adjacent normal cells. Pathological diagnosis and treatment assessment is based on the subtype that the breast cancer tumor cells fall into based on 1) clinical staging, 2) histology, 3) biomarker expression and 4)molecular profiling by gene expression^35–37^. This study demonstrates the TME in breast cancer based on a variety of spatial data and technologies is very consistent with pathological anatomical features in relation to stemness. We present for the first time a ML model that can predict stemness while preserving the spatial context of cell interactions within the tissue. This model highlights the importance of understanding where in the tumor, more aggressive, stem-like cells are situated and the TME that surrounds them. This model preserves single cell identities and can incorporate existing single cell data to inform which cell types are more stem-like, highlighting the adaptability to existing single cell datasets and the integration into emerging spatial omic datasets. This model is also agnostic to which single cell or spatial technology is used to generate the input datasets.

Because the TME is a critical aspect of breast cancer management, this study lends to the importance of harnessing cell identities, composition and interactions from single cell and spatial omics data to better our understanding of clinical outcomes and treatment. The ease of use and speed of this model makes it highly attractive to generate stemness prediction and discover pathogenicity of the tumor when confronted with analyzing millions of cells at a time. Overall, ML-based methods can provide a powerful toolset for scRNAseq and spatial omics analysis, which can help accelerate our understanding of cancer stemness and its clinical relevance.

## Figure legends

**Supplemental Figure 1** (A) Stemness on bootstrapped high and poor quality breast cancer samples on dataset 1 (i,ii) and dataset 2 (iii,iv). Each figure represents one sample and the box plot indicates the stemness distribution of a subset of cells that grows in number to the right until the total cells for that given sample. (B) Rank barplot of number of cells by patient samples for dataset 1(i) and dataset 2 (ii).

**Supplemental Figure 2** Gene expression analysis of stemness model. Volcano plots of top stemness A) cancer cycling cells from dataset1 (i) and dataset 2 (ii) respectively. Gradient depicts negative logFC (blue) to positive logFC (red). Y axis represents the statistical significance and X axis the different percentage of cells expressing a given gene. Positive different percentage means that top stemness cells have more cells expressing that gene. B) GO enrichment analysis of the top regulated genes in cancer cycling cells for dataset 1 (red) and dataset 2 (blue). The number of DEG is plotted on the X axis and GO categories the genes are represented in are plotted on the y axis. C) Volcano plots of top stemness cycling T cells from dataset1 (i) and dataset 2 (ii) respectively. Gradient depicts negative logFC (blue) to positive logFC (red). D) GO enrichment analysis comparison from the number of up regulated genes in cycling t-cells dataset 1 (red) and dataset 2 (blue). E) Gene similarity score of upregulated genes from cancer basal high stemness cells vs cluster markers genes. On the left, Dataset 1 high stemness genes are compared to gene markers from Dataset 2. On the right, Dataset 2 high stemness genes are compared to gene markers from Dataset 1. Gradient depicts less gene similarity (blue) to greater gene similarity (red).

**Supplemental Figure 3** (A) Heatmap of gene similarity analysis of cluster markers in dataset 1 compared to dataset 2 for cancer basal DEGs and (B) heatmap of gene similarity analysis of cluster markers in dataset 2 compared to dataset 1 for the same cell type.

**Supplemental Figure 4** QC Metrics of Akoya PhenoCycler and Nanostring CosMx samples. A) Violin plot of fluorescence intensity signal (nCount_Akoya) showing the number of proteins detected per cell (nFeature_Akoya). B) Scatter plot of number of fluorescence intensity (x-axis) by number of proteins detected per cell (y-axis) and the title shows the pearson correlation value. C) Violin plot of fluorescence intensity signal (nCount_Nanostring) showing the number of proteins detected per cell(Feature_Nanostring) D) Scatter plot of number of fluorescence intensity (x-axis) by number of proteins detected per cell (y-axis) and the title shows the pearson correlation value.

## Notes

### Competing Interest Statement

The authors have declared no competing interest.

