## Supplemental Figures for "A Machine Learning One-Class Logistic Regression Model to Predict Stemness in Single Cell Transcriptomics and Spatial Omics Datasets"

A)

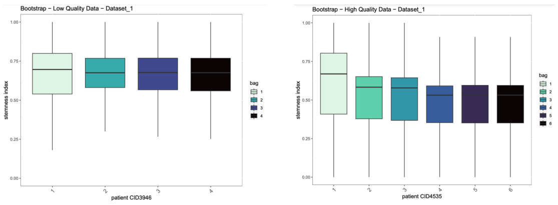


B)

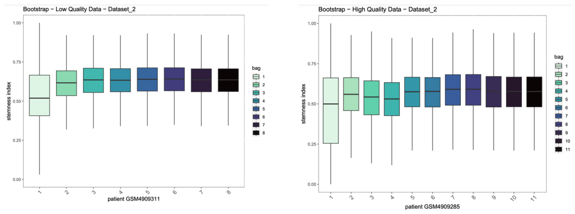


C)

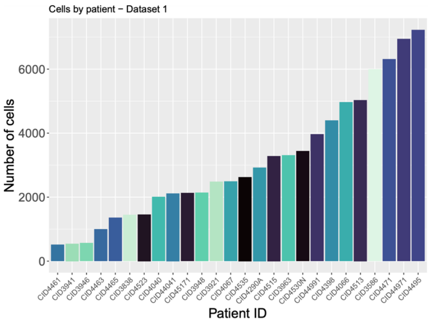


D)

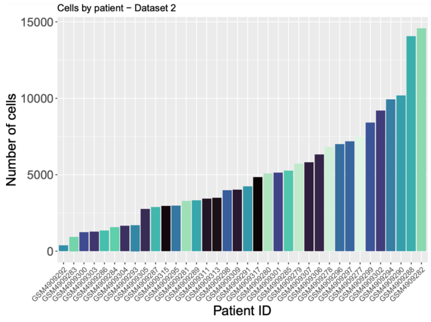


### Supp. Figure 1

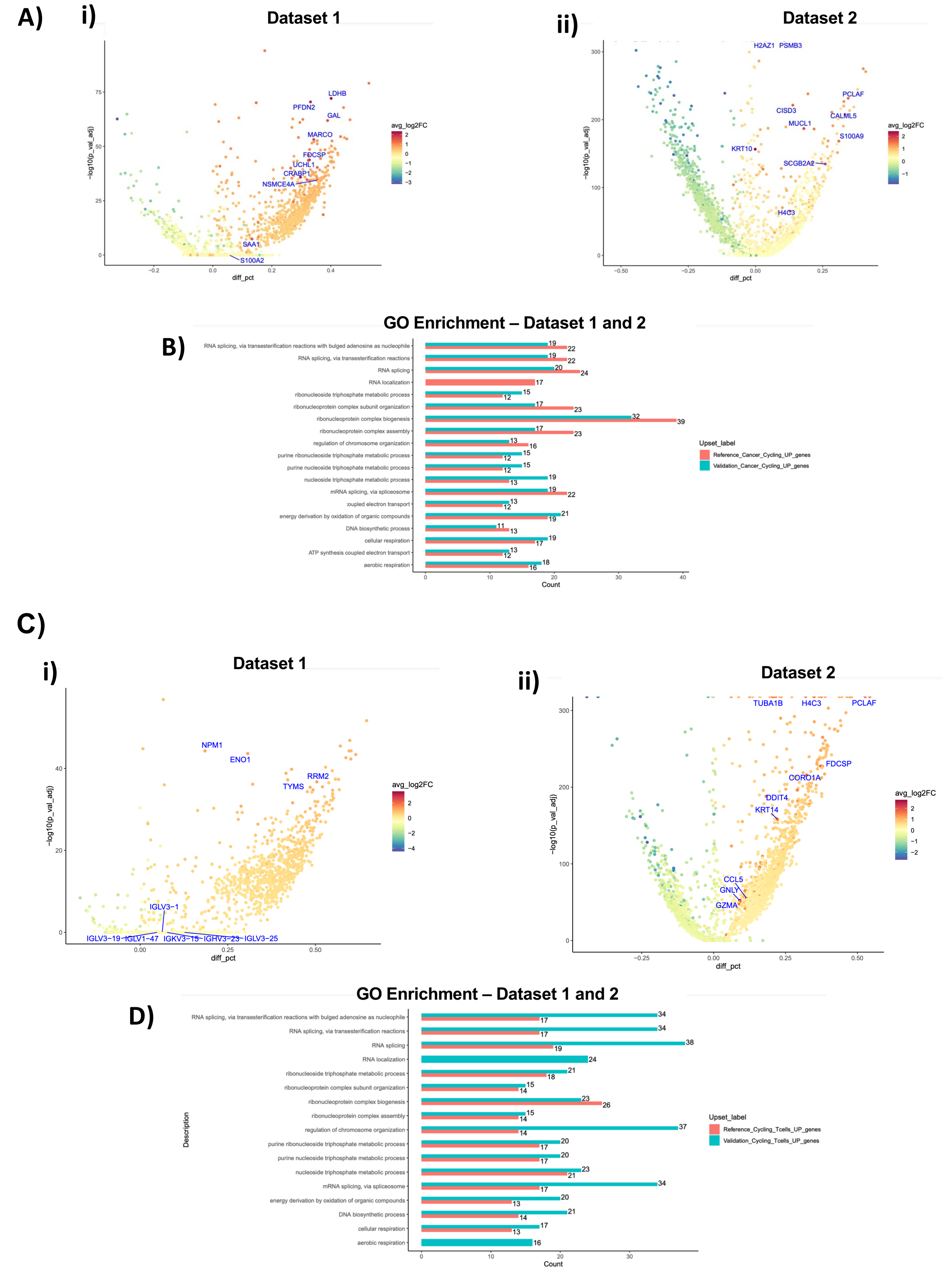


Supp. Figure 2

A)

Cancer basal - Dataset 1 vs Dataset 2 – Cluster Markers gene similarity

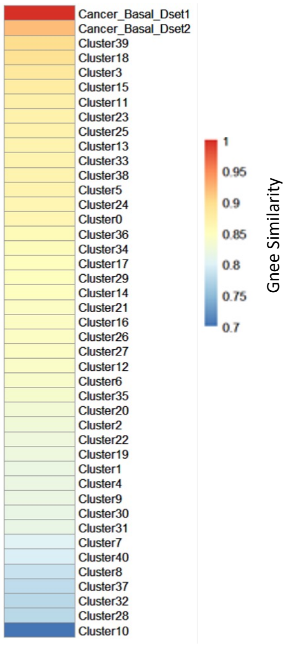


B)

Cancer basal - Dataset 2 vs Dataset 1 – Cluster Markers gene similarity

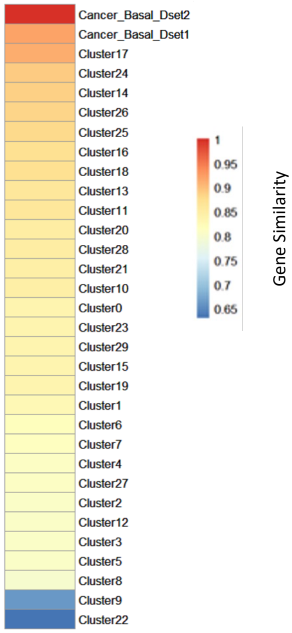


### Supp. Figure 3

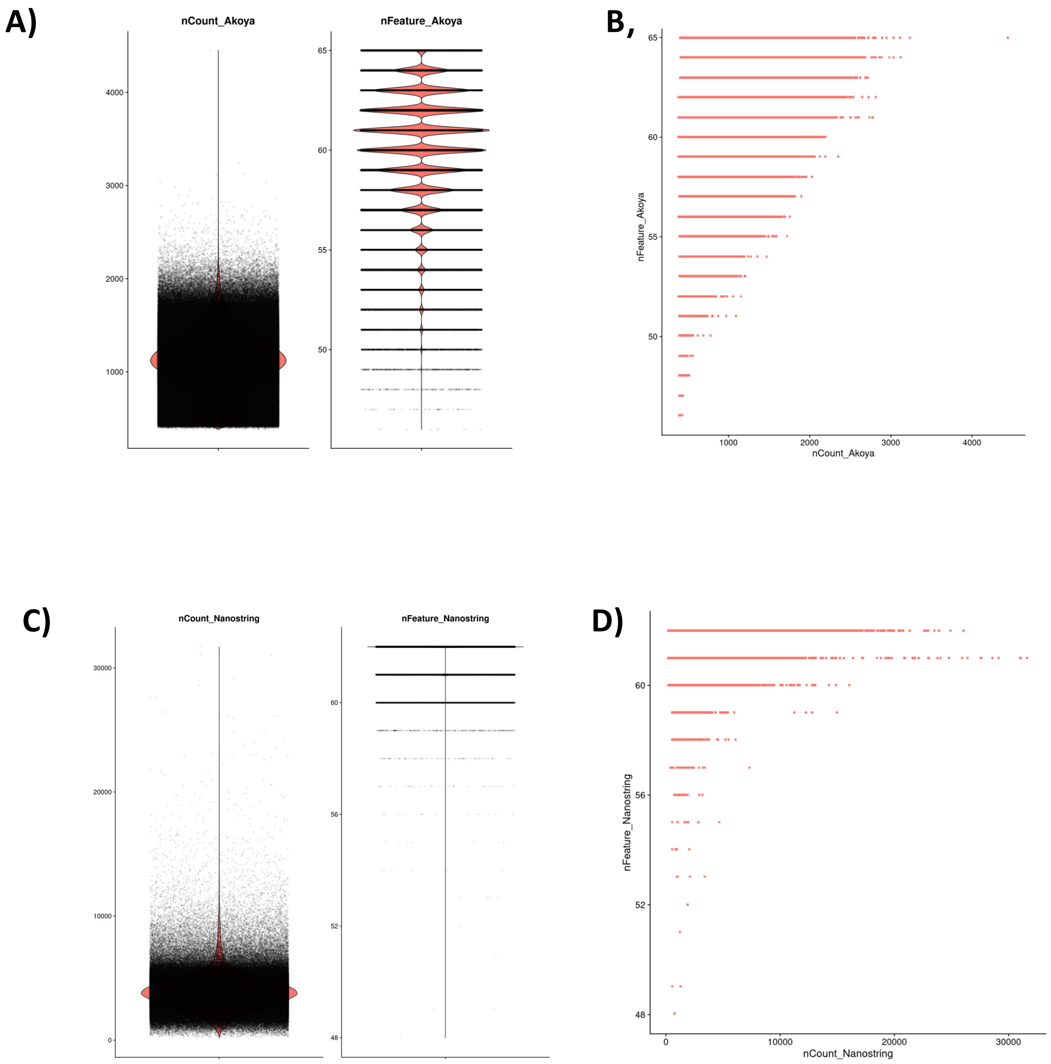


### Supp. Figure 4
